# Control of myogenesis by the E3 ubiquitin ligase CUL3-BTBD9

**DOI:** 10.1101/2025.05.15.654347

**Authors:** Chris Padovani, Fernando Rodriguez-Perez, Monica Tsai, Jay Xiong, Angela Pogson, Brenda Martinez, Michael Rape

**Affiliations:** Department of Molecular and Cell Biology, University of California at Berkeley, Berkeley, CA 94720 USA; Howard Hughes Medical Institute, University of California at Berkeley, Berkeley, CA 94720, USA; California Institute for Quantitative Biosciences (QB3), University of California at Berkeley, Berkeley, CA 94720, USA; Eikon, Hayward, CA 94545, USA; Stanford University, Stanford, CA 94305, USA

**Keywords:** ubiquitin, BTBD9, CUL3, myogenesis, CAV1, KCTD10, insulin

## Abstract

Metazoan development requires that cells adopt specific identities at the right time and place within an embryo. Central to the success of this process are posttranslational modifications that control the activity, stability or localization of crucial transducers of differentiation signals. During muscle development, modification of proteins with ubiquitin is known to play an important role, but the enzymatic machinery of ubiquitylation that drives myogenesis remains incompletely understood. Here, we identify CUL3^BTBD9^ as an E3 ubiquitin ligase that is essential for myogenesis *in vitro*. CUL3^BTBD9^ binds and ubiquitylates CAV1, the central component of caveolae that modulate insulin signaling during muscle formation. CUL3^BTBD9^ and CAV1 are required for insulin-dependent activation of the AKT kinase in myoblasts, thereby safeguarding the ability of muscle precursors to respond to insulin signals. Together, this work identifies CUL3^BTBD9^ as a regulator of myogenesis that acts by modulating plasma-membrane localized events critical for cell fate specification.

## Introduction

Metazoan development requires that stem cells accurately interpret environmental signals to elicit differentiation events that produce specific cell types at the right time and place within an embryo. The exit of stem cells from their self-renewal program and the concomitant initiation of differentiation is accordingly tightly controlled. Critical to this regulation are posttranslational modifications that modulate the activity, localization or stability of key transducers of differentiation signals. Identifying enzymes that catalyze such modifications has provided important insight into components and regulators of signaling pathways that are responsible for cell differentiation, and mutations in such enzymes have frequently been linked to developmental diseases ^1–3^.

Posttranslational modification with ubiquitin is emerging as an important posttranslational modification guiding cell fate specification ^1^. Ubiquitylation requires a cascade of E1 ubiquitin activating enzymes, E2 conjugating enzymes, and E3 ligases ^4–6^. Together, these enzymes decorate proteins with single ubiquitin molecules, known as monoubiquitylation, or with polymeric ubiquitin chains that can adopt homotypic or complex heterotypic topologies ^5, 7–10^. The type of modification determines its consequences: while monoubiquitylation predominantly regulates protein localization or interactions, polymeric chains often elicit protein degradation by the proteasome or autophagy. Both mono- and polyubiquitylation play important roles during cell fate specification. For example, monoubiquitylation regulates chromatin architecture to allow transcription factors to initiate the gene expression programs driving cell differentiation ^11, 12^. Conversely, modification of transcription factors with ubiquitin polymers induces their degradation and thereby ensures that stem cells and their differentiating counterparts remain susceptible to the environmental signals that guide the differentiation program ^13, 14^. The substrate specificity of ubiquitylation is determined by E3 ligases, whose identification has accordingly become an important step towards revealing regulatory networks underlying successful cell differentiation and development.

The central role of ubiquitylation for cell fate specification has been illustrated during myogenesis, a developmental program that produces multinucleate myotubes as the characteristic building blocks of muscle ^15^. Some E3 ligases have been found to play critical roles during muscle formation or homeostasis. For example, NEDD4 controls the PAX7 transcription factor that is crucial for proper execution of gene programs in muscle stem cells ^16^. The E3 CUL3^KCTD10^ monoubiquitylates actin-bundling proteins at the plasma membrane to control the cell fusion events that enable myotube formation ^17^. CUL2^FEM1B^ acts as a component of the reductive stress response that allows myoblasts to adjust their mitochondrial electron transport chain to requirements of myotube formation ^18–20^. TRIM63 targets structural proteins in myofibrils for degradation and thereby accelerates muscle degeneration during sarcopenia ^21–24^, and SCF^FBXO32^ impacts myotube integrity by ubiquitylating a translation initiation factor as well as intermediate filament proteins ^25, 26^. Recent genetic screens identified additional E3 ligases with likely roles in myogenesis ^19^, but cellular targets and regulation of these enzymes are still unknown. Important functions of ubiquitylation in myogenesis therefore remain to be discovered.

Here, we identify CUL3^BTBD9^ as an E3 ligase that is essential for myogenesis. CUL3^BTBD9^ binds and ubiquitylates CAV1, the key component of caveolae that modulate many plasma membrane-localized signaling events ^27, 28^. CUL3^BTBD9^ recognizes the scaffold domain of CAV1 that is known to mediate protein interactions required for signal transduction, leading to modification with non-proteolytic ubiquitin oligomers. CUL3^BTBD9^ and CAV1 are both required for insulin-dependent activation of the AKT kinase, safeguarding the ability of myoblasts to respond to insulin as a critical developmental signal. Our work therefore identifies an E3 ligase, CUL3^BTBD9^, that promotes myogenesis by controlling signaling events at the plasma membrane.

## Results

### BTBD9 is required for myoblast differentiation

We recently performed genetic screens to find E3 ligases required for myoblast differentiation, an approach that allowed us to discover the reductive stress response built around CUL2^FEM1B^ ^19^. As another top hit of these screens, depletion of the E3 ligase subunit BTBD9 very strongly inhibited myoblast differentiation (**Figure S1A**). We now revisited these findings with independent siRNAs and found that reducing BTBD9 levels abolished myogenesis, as detected by a lack of multinucleate myotubes positive for the late differentiation marker myosin heavy chain (MyHC; **Figure S1B**). Differentiation was restored upon expressing siRNA-resistant BTBD9, showing that effects of siRNAs were due to the on-target loss of BTBD9 (**Figure S1C**).

Using CRISPR/Cas9-mediated genome engineering, we inactivated endogenous *BTBD9* in C2C12 myoblasts. Mirroring the consequences of transient depletion, *ΔBTBD9* myoblasts were unable to induce myotube formation, as seen by immunofluorescence microscopy detecting early or late differentiation markers, the transcription factor myogenin (MyoG) or MyHC, respectively (**Figure 1A, B**). The defect in cell fate specification of *ΔBTBD9* myoblasts was also noticeable by Western blotting (**Figure 1C**). These experiments thus showed that the E3 ligase subunit BTBD9 is essential for C2C12 myogenesis *in vitro*.

**Figure 1:**
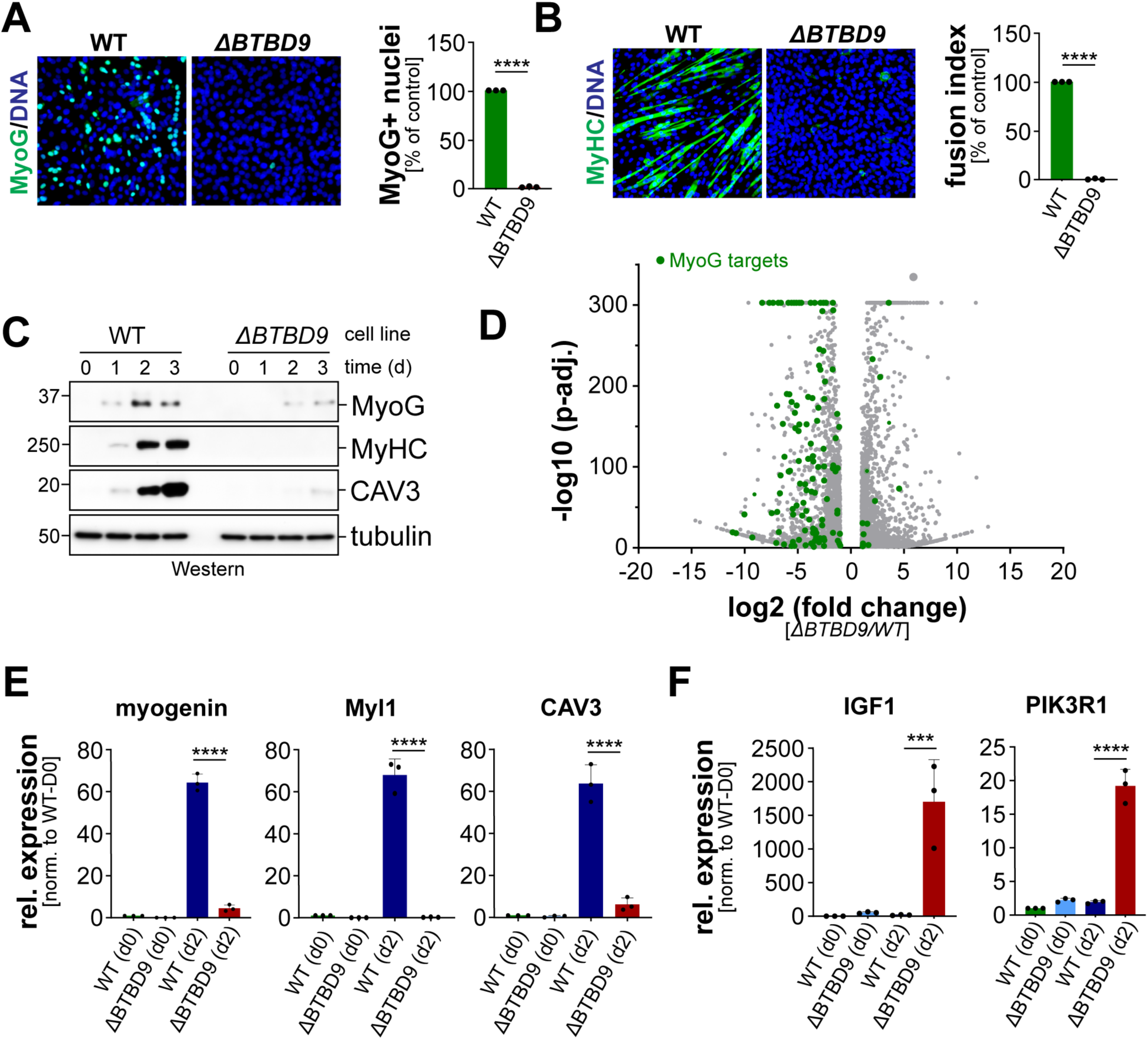
BTBD9 is required for *in vitro* myogenesis. **A.** Wildtype or *ΔBTBD9* C2C12 myoblasts were transferred to differentiation medium. 2d later, expression of the early differentiation marker myogenin was detected by immunofluorescence microscopy (*green:* myogenin, MyoG; *blue:* DNA). Quantification of three independent experiments in shown on the right. **B.** Wildtype or *ΔBTBD9* C2C12 myoblasts were transferred to differentiation medium. 2d later, expression of the late differentiation marker MyHC was detected by immunofluorescence microscopy (*green:* MyHC; *blue:* DNA). Quantification of three independent experiments in shown on the right. **C.** Differentiation of wildtype or *ΔBTBD9* C2C12 myoblasts was followed by Western blotting, detecting the indicated differentiation markers with specific antibodies. **D.** Wildtype or *ΔBTBD9* C2C12 myoblasts were transferred to differentiation medium. 2d later, gene expression was analyzed by RNA-sequencing (*green:* targets of the muscle transcription factor myogenin, MyoG). **E.** Analysis of expression of myogenesis markers using qRT-PCR, in WT or *ΔBTBD9* C2C12 myoblasts either prior to initiation of differentiation (d0) or two days after initiation of differentiation (d2). **F.** Analysis of expression of IGF1 or PI3KR1 in WT or *ΔBTBD9* C2C12 myoblasts either prior to initiation of differentiation (d0) or two days after initiation of differentiation (d2).

To understand how BTBD9 drives myogenesis, we subjected WT or *ΔBTBD9* myoblasts to a differentiation protocol that is characterized by growth factor withdrawal. After two days, when wildtype cells have started to differentiate, we analyzed gene expression by RNA-sequencing (**Figure 1D**). These experiments revealed that the absence of *BTBD9* caused an early disruption of myogenesis, as targets of the transcription factors MyoD and myogenin were strongly reduced in *ΔBTBD9* compared to WT cells (**Figure 1D; Figure S2A**). We confirmed the reduced expression of myogenesis markers by qRT-PCR (**Figure 1E**). Conversely, we noted that *ΔBTBD9* cells increased the expression of mRNAs encoding proteins linked to growth factor and insulin signaling, such as insulin-like growth factor 1 (IGF1) and phosphoinositol-3-kinase (PI3K), an observation that we also validated by qRT-PCR (**Figure 1F**). Consistent with these results, GO term analyses showed that the PI3K/AKT kinase pathway that acts downstream of insulin signaling was most strongly affected by *BTBD9* deletion (**Figure S2B**).

We conclude that BTBD9 is required for the initiation of myogenesis *in vitro*. BTBD9 is a substrate adaptor of CUL3, which itself is known to play important roles during myoblast differentiation ^19, 29, 30^. BTBD9 possesses an N-terminal BTB domain that binds CUL3, a BACK domain, and two discoidin domains that in other proteins mediate important interactions ^31, 32^ (**Figure 2A**). Polymorphisms in intronic regions of the *BTBD9* gene are associated with Restless Leg and Tourette syndromes ^33–35^, and deletion of *BTBD9* in *Drosophila* or mice altered synaptic transmission through changing the expression of regulators or targets of endocytosis ^36, 37^. The strong effects of *BTBD9* deletion on myogenesis and the multiple links between BTBD9 and development and disease prompted us to investigate how CUL3^BTBD9^ controls myogenesis.

**Figure 2:**
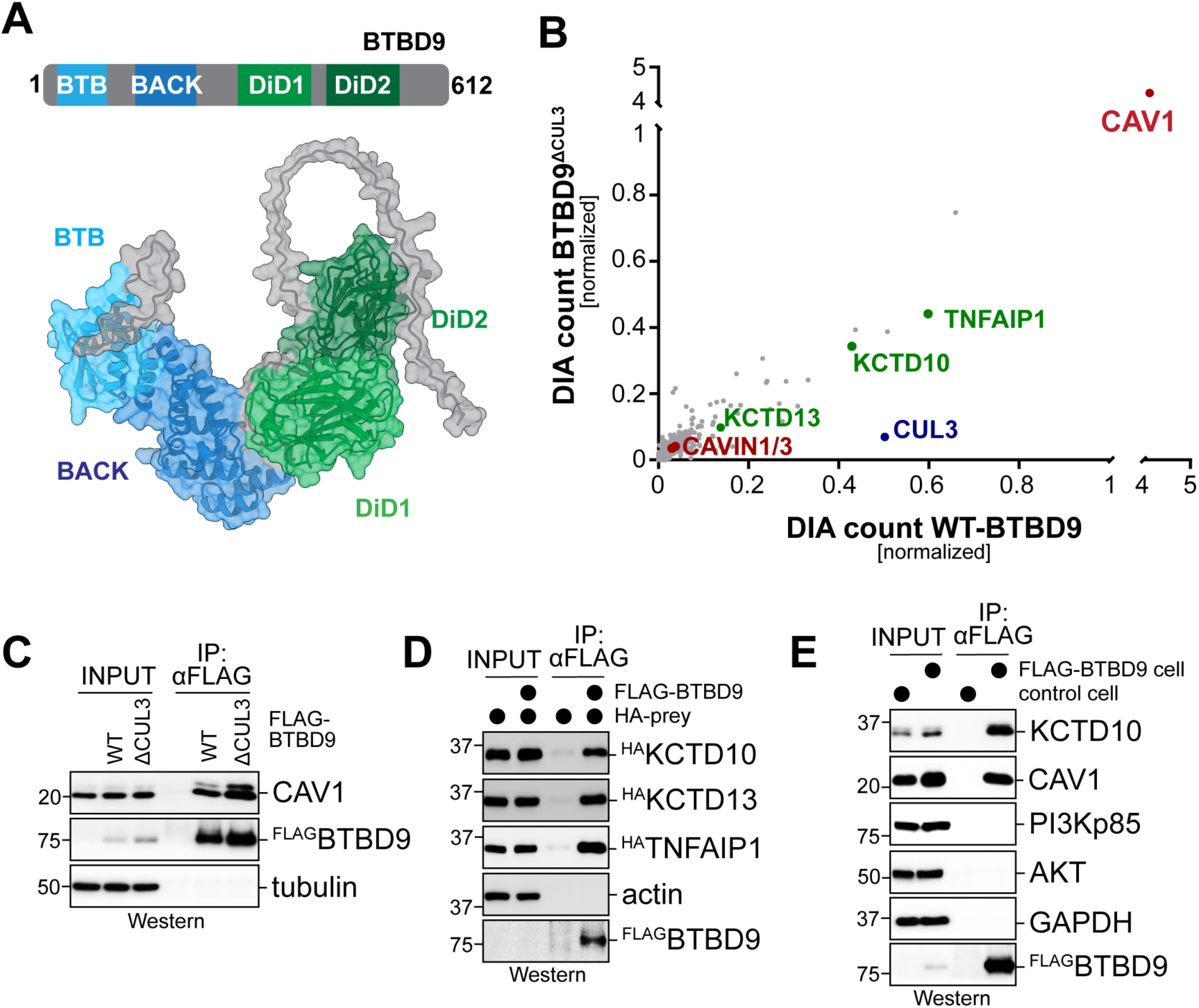
BTBD9 binds CAV1. **A.** *Upper panel:* Domain structure of BTBD9 (DiD: discoidin domain). *Lower panel:* AlphaFold model of BTBD9. **B.** Data-independent acquisition mass spectrometry analysis of affinity-purifications of wildtype BTBD9 (x-axis) and BTBD9^ΔCUL3^ (y-axis). **C.** ^FLAG^BTBD9 or ^FLAG^BTBD9^ΔCUL3^ were affinity-purified from C2C12 myoblasts and endogenous co-precipitating CAV1 was detected by Western blotting. **D.** ^FLAG^BTBD9 was co-expressed in C2C12 myoblasts with indicated HA-tagged proteins (‘prey’), and co-purifying proteins were detected by Western blotting. **E.** Control C2C12 myoblasts of *BTBD9::FLAG* myoblasts (endogenous *BTBD9* loci were tagged with FLAG) were subjected to αFLAG affinity-purification, and co-precipitating endogenous proteins were detected by Western blotting.

### CUL3^BTBD9^ binds KCTD10 and CAV1

We started by identifying BTBD9 interactors as potential targets or regulators of CUL3^BTBD9^. We expressed in C2C12 myoblasts either wildtype ^FLAG^BTBD9 or ^FLAG^BTBD9^M69A/R70A/E71A^ (referred to as BTBD9^ΔCUL3^), a variant that is defective in CUL3-binding and cannot ubiquitylate substrates. Especially if ubiquitylation results in degradation, the delayed product release from BTBD9^ΔCUL3^ should facilitate substrate identification by mass spectrometry. We immunoprecipitated BTBD9 and determined binding partners by data-independent acquisition mass spectrometry. BTBD9, but much less BTBD9^ΔCUL3^, engaged CUL3, as expected for a substrate adaptor of a CUL3 E3 ligase (**Figure 2B**). In addition, we found that BTBD9 abundantly associated with CAV1, the major constituent of caveolae that regulate insulin signaling and muscle stem cell differentiation ^27, 28^. BTBD9 affinity-purifications also contained the CAVIN1 and CAVIN3 proteins that bind CAV1 at caveolae. In addition, BTBD9 bound three CUL3 adaptors that are related to each other, KCTD10, KCTD13 and TNFAIP1, and KCTD10 had previously been shown to mediate cell fusion during myogenesis ^17^. BTBD9^ΔCUL3^ recognized CAV1, KCTD10, KCTD13, and TNFAIP1 with similar efficiency as wild-type BTBD9, which implied that these interactors are either substrates of non-proteolytic ubiquitylation or regulators of CUL3^BTBD9^.

Analysis of BTBD9 affinity-purifications by Western blotting confirmed that BTBD9 strongly associated with endogenous CAV1 (**Figure 2C**), and it also engaged heterologously expressed KCTD10 and TNFAIP1 (**Figure 2D**). To assess these interactions without overexpression, we fused FLAG epitopes to *BTBD9* loci in C2C12 myoblasts. Following immunoprecipitation of endogenous BTBD9, we found that it efficiently interacted with CAV1 and KCTD10 (**Figure 2E**). BTBD9 therefore strongly interacts with two proteins, CAV1 and KCTD10, that each act at the plasma membrane and have been found to regulate myoblast differentiation. However, neither CAV1 (**Figure S3A**) nor KCTD10 (ref. ^17^) are essential for myogenesis, suggesting that CUL3^BTBD9^ regulates additional, still unknown, factors to exert its control over muscle stem cell differentiation.

### BTBD9 uses its second discoidin domain to engage CAV1 and KCTD10

By deleting specific domains in BTBD9, we found that its second discoidin domain was required for recognition of both KCTD10 and CAV1 (**Figure 3A**). Discoidin domains are interaction modules that mediate complex formation at the plasma membrane and help transmit extracellular signals into the cytoplasm ^32^. AlphaFold suggested with high confidence that the second discoidin domain of BTBD9 engages a predicted thioredoxin-like domain in KCTD10 (**Figure 3B**). Specifically, three residues in BTBD9 (N546, E547, V548) dock into a hydrophobic pocket on the surface of KCTD10. Mutation of these residues strongly impaired the binding of BTBD9 to KCTD10 (**Figure 3C**). Immunoprecipitation coupled to mass spectrometry confirmed that the NEV motif in the second discoidin domain of BTBD9 was required for efficient binding to KCTD10, but not CUL3 (**Figure 3D**). We will refer to a mutant in the NEV-motif of BTBD9 as BTBD9^DiD2-M^.

**Figure 3:**
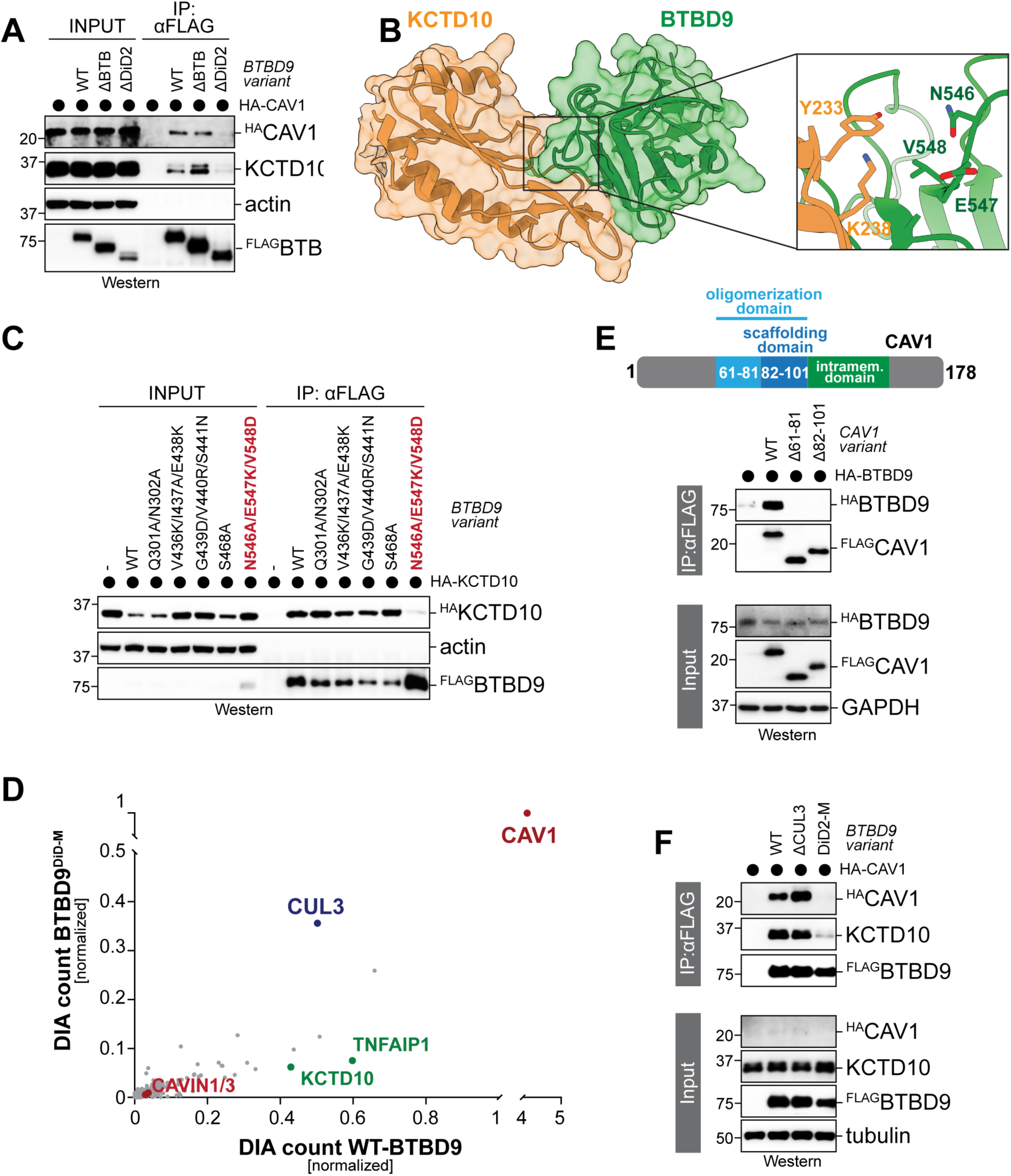
The second discoidin domain of BTBD9 recognizes CAV1. **A.** FLAG-tagged wild-type BTBD9 or variants with a mutant CUL3-binding interface (ΔCUL3) or with a deletion removing the second discoidin domain (ΔDiD2) were co-expressed with ^HA^CAV1. Following α-FLAG affinity-purification, co-precipitating proteins were detected by Western blotting. **B.** AlphaFold model (iPTM 0.72) of a complex between a thioredoxin-like domain in KCTD10 and the second discoidin domain in BTBD9. **C.** BTBD9 variants with indicated mutations were affinity-purified, and co-precipitating ^HA^KCTD10 was detected by Western blotting. **D.** Mass spectrometry analyses of immunoprecipitated BTBD9 or BTBD9 with mutations in three critical residues required for substrate engagement (BTBD9^DiD2-M^). **E.** The oligomerization and scaffolding domain of CAV1 is required for recognition by BTBD9. FLAG-tagged CAV1 variants were affinity-purified and co-precipitating BTBD9 was detected by Western. **F.** The second discoidin domain of BTBD9 is required for CAV1 recognition. The indicated BTBD9 variants were affinity-purified from C2C12 myoblasts, and co-precipitating proteins were detected by Western blotting.

As AlphaFold could not predict the binding interface between BTBD9 and CAV1, we proceeded by generating CAV1 deletion mutants to identify domains needed for engaging BTBD9. These experiments revealed that deletions in the oligomerization and scaffold domain of CAV1, a hotspot for interactions with regulators of signal transduction ^28^, impaired binding to BTBD9 (**Figure 3E**). As seen with KCTD10, mutation of the NEV motif in BTBD9 disrupted the recognition of CAV1 (**Figure 3F**), which was also observed for endogenous CAV1 by affinity-purification and mass spectrometry (**Figure 3D**). Despite the shared requirement for the NEV motif in BTBD9, sequential affinity-purifications showed that BTBD9-CAV1 complexes contained KCTD10 (**Figure S3B**), suggesting that both BTBD9-interactors are present in overlapping signaling complexes. We conclude that KCTD10 and CAV1 are interactors of BTBD9 that are specifically recognized by the second discoidin domain of BTBD9.

### CUL3^BTBD9^ ubiquitylates CAV1

We noted that some CAV1 bound to wildtype BTBD9, but not to BTBD9^ΔCUL3^, was modified consistent with monoubiquitylation by CUL3^BTBD9^ (**Figure 4A**). To test whether CUL3^BTBD9^ ubiquitylates CAV1, we restored *ΔBTBD9* cells with either wildtype-BTBD9 or CUL3-/CAV1-binding variants. Following expression of HIS6-tagged ubiquitin, we enriched conjugates from cell lysates by denaturing purification. We found that WT-BTBD9 strongly induced modification of CAV1 with up to three ubiquitin molecules (**Figure 4B**). Underscoring the specificity of this reaction, neither the BTBD9 variant defective in CUL3-binding (BTBD9^ΔCUL3^) nor the variant that cannot engage CAV1 (BTBD9^DiD2-M^) was able to promote CAV1 modification. Ubiquitylation of CAV1 was not affected by concomitant overexpression of KCTD10 (**Figure S4A**). While KCTD10 was also modified in cells, BTBD9 deletion did not diminish this modification, nor was its ubiquitylation status affected by mutations in the CUL3- or KCTD10-binding sites in overexpressed BTBD9 (**Figure S4B**). Thus, CUL3^BTBD9^ targets CAV1 for ubiquitylation, while modification of KCTD10 might require additional factors. Alternatively, KCTD10 might interact with BTBD9 in a role that is different from that of a substrate.

**Figure 4:**
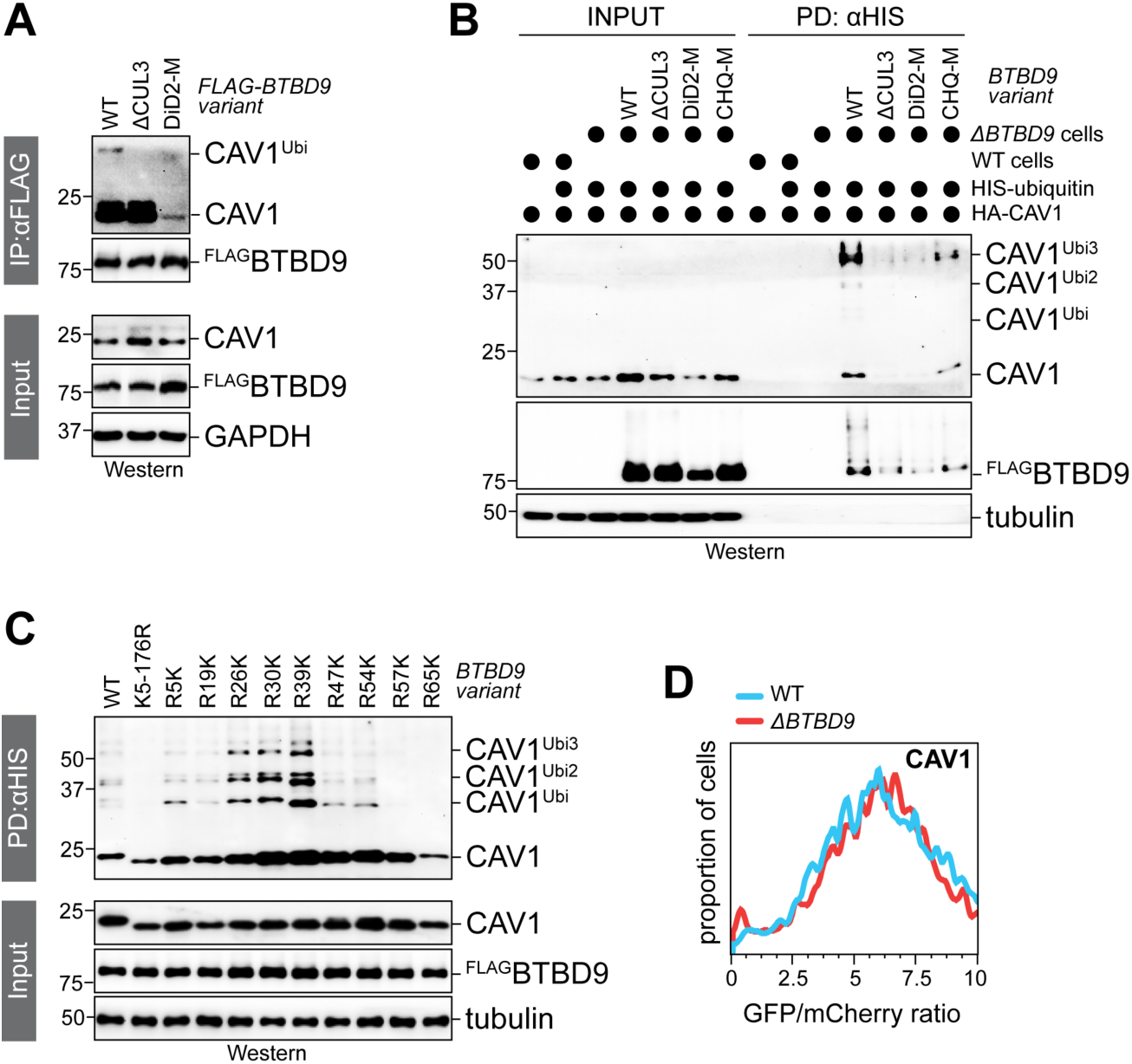
BTBD9 promotes non-proteolytic ubiquitylation of CAV1. **A.** Indicated ^FLAG^BTBD9 variants were affinity-purified and co-precipitating CAV1 was detected by Western blotting. Only affinity-purification of WT-BTBD9, but not a mutant defective in catalyzing ubiquitylation (ΔCUL3) resulted in co-precipitation of a modified version of CAV1, with a molecular weight expected for monoubiquitylated CAV1. **B.** Denaturing NiNTA-purification of His-ubiquitin conjugates from either WT or *ΔBTBD9* myoblasts shows that overexpression of BTBD9 resulted in modification of CAV1 with up to three ubiquitin molecules, as detected by Western blotting. **C.** Denaturing NiNTA-purification of His-ubiquitin conjugates from cells expressing single-Lys variants of CAV1 reveals that BTBD9 preferentially promotes modification of K26, K30, and K39 in CAV1 with short oligomers containing up to three ubiquitin subunits. **D.** Degradation reporter assay, as described ^18^, shows that deletion of *BTBD9* did not alter the stability of CAV1.

As the consequences of modification with one or few ubiquitin molecules can depend on the site of ubiquitin attachment ^38^, we next established CAV1 variants that possessed only a single Lys residue within the amino-terminal 176 residues. Using denaturing purification of ubiquitin conjugates, we found that CUL3^BTBD9^ most effectively modified K39 in CAV1 (**Figure 4C**). CUL3^BTBD9^ could also target the neighboring K26 and K30, and it modified each single-Lys variant with up to 3-4 ubiquitin subunits. These experiments showed that CUL3^BTBD9^ forms ubiquitin oligomers on CAV1 residues near the scaffolding domains that recruit BTBD9. Unfortunately, we were unable to reconstitute this reaction *in vitro*, which prevented us from determining if CAV1 is modified with ubiquitin oligomers of specific topology. Given that AlphaFold was also unable to predict the binding interface between CAV1 and BTBD9, it is possible that additional factors or posttranslational modifications are required for successful ubiquitylation of CAV1 by CUL3^BTBD9^.

While mono- or oligoubiquitylation can alter protein function, degradation typically requires polymeric chains ^5,7^. To further probe whether CUL3^BTBD9^ modifies CAV1 with low molecular weight conjugates, we assessed CAV1 stability in *ΔBTBD9* cells using stability reporters that can monitored by flow cytometry, as described ^13, 19, 20, 39–43^. As expected from the ubiquitin oligomers that we found to be attached to CAV1, deletion of *BTBD9* did not stabilize CAV1 (**Figure 4D**). These findings indicated that CUL3^BTBD9^ modifies CAV1 with ubiquitin mono- or oligomers that serve non-proteolytic roles.

### BTBD9 regulates CAV1 expression

As CUL3^BTBD9^ modifies CAV1 with non-proteolytic ubiquitin conjugates, we next asked if the E3 ligase regulates CAV1 function or regulation. Mass spectrometry analyses of CAV1 immunoprecipitates from either wildtype or *ΔBTBD9* cells showed that many CAV1 interactions were altered by the absence of BTBD9, which included proteins with roles in myogenesis (**Figure 5A**). Given the complexity of these changes, it was difficult to pinpoint a single binder whose function was modulated by CUL3^BTBD9^-dependent CAV1 ubiquitylation. In addition, we found by Western blotting or immunofluorescence microscopy that loss of *BTBD9* caused a strong increase in CAV1 levels, which was particularly noticeable if myoblasts had been transferred to differentiation conditions low in growth factors (**Figure 5B, C**). These observations suggested that CUL3^BTBD9^ might also regulate, either directly or indirectly, the expression of CAV1.

**Figure 5:**
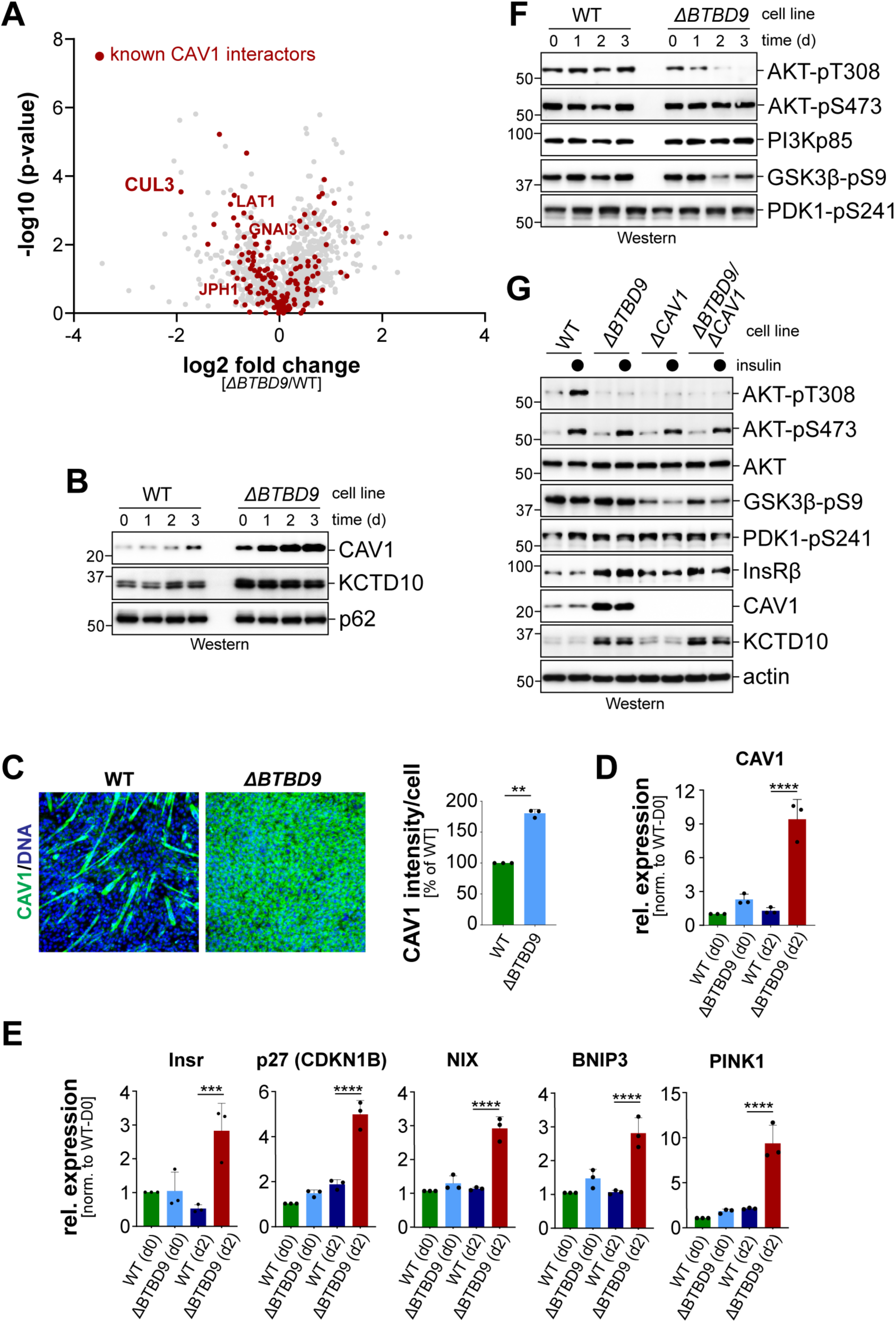
Deletion of *BTBD9* impacts CAV1 expression and AKT activation. **A.** Mass spectrometry analyses of immunoprecipitated ^FLAG^CAV1 from either WT or *ΔBTBD9* myoblasts. Known CAV1 interactors are shown in red. **B.** WT or *ΔBTBD9* C2C12 myoblasts were transferred to differentiation medium, and the indicated proteins were detected over time by Western blotting. **C.** WT or *ΔBTBD9* C2C12 myoblasts were analyzed for CAV1 expression by immunofluorescence microscopy against the endogenous proteins. Quantification of three independent experiments is shown on the right. **D.** Analysis of expression of CAV1 mRNA using qRT-PCR, in WT or *ΔBTBD9* C2C12 myoblasts either prior to initiation of differentiation (d0) or two days after initiation of differentiation (d2). **E.** Analysis of expression of various FOXO target genes using qRT-PCR, in WT or *ΔBTBD9* C2C12 myoblasts either prior to initiation of differentiation (d0) or two days after initiation of differentiation (d2). **F.** WT or *ΔBTBD9* C2C12 myoblasts were transferred to differentiation medium, and the indicated components of the PI3K-AKT signaling pathway were detected over time by Western blotting. **G.** The indicated C2C12 myoblasts lines were subjected to brief serum starvation and re-stimulated with insulin. Protein expression was analyzed by Western blotting.

Consistent with the latter notion, we found by qRT-PCR that CAV1 mRNA levels were increased in *ΔBTBD9* myoblasts that were subjected to differentiation by partial growth factor withdrawal (**Figure 5D**). In previous work, CAV1 expression had been shown to be driven by FOXO transcription factors that are inhibited by AKT ^44^. We analyzed expression of additional FOXO target genes and found that their transcription was increased upon *BTBD9* deletion in a manner similar to CAV1 (**Figure 5E**). *ΔBTBD9* myoblasts induced these genes most prominently after being transferred to differentiation medium low in growth factors. CUL3^BTBD9^ therefore regulates the expression of FOXO target genes that include its ubiquitylation target, CAV1.

Having seen the effects of *BTBD9* deletion on the expression of FOXO target genes, we asked if loss of CUL3^BTBD9^ led to a dysregulation of AKT. Note that our initial RNA sequencing experiments had indicated dysregulation of PI3K/AKT signaling in *ΔΒΤΒD9* myoblasts (**Figure S2B**). The activation status of AKT downstream of insulin/PI3K signaling can be followed by monitoring phosphorylation of its T308 residue dependent on the upstream kinase PDK ^45^. We found that *BTBD9* deletion led to a gradual loss of AKT-T308 phosphorylation, if cells were transferred to differentiation conditions characterized by partial growth factor withdrawal (**Figure 5F**). We next repeated this experiment using myoblasts that were acutely stimulated with insulin after short periods of starvation. While insulin induced AKT phosphorylation at T308 in wildtype myoblasts, this signaling was blocked if *BTBD9* had been inactivated (**Figure 5G**). Consistent with CUL3^BTBD9^ and CAV1 acting in the same pathway, *ΔCAV1* myoblasts also showed a strong reduction in AKT acti-vation by insulin stimulation. Phosphorylation of AKT at Ser473 through mTORC2 was only mildly impacted by the absence of *BTBD9* or *CAV1*. Together, these observations implied that CUL3^BTBD9^ and CAV1 collaborate to allow for activation of the AKT kinase that plays important roles in myoblast gene expression and differentiation ^46^.

### BTBD9 regulates insulin dynamics and signaling

AKT is activated upon binding of insulin or insulin-like growth factor 1 (IGF1) to their receptors at the plasma membrane. Using immunofluorescence microscopy of endogenous BTBD9, we found that BTBD9 was enriched at the plasma membrane (**Figure 6A**), where its localization partially overlapped with that of IGF1 receptor (**Figure 6A**). Interestingly, CUL3-inhibition by MLN4924 or deletion of *CAV1* resulted in an enrichment of IGF1 receptor in membrane domains that also contained BTBD9 (**Figure 6A**). These results implied that CUL3^BTBD9^ acts in a CAV1-dependent manner to restrict the plasma membrane accumulation of IGF1 receptor.

**Figure 6:**
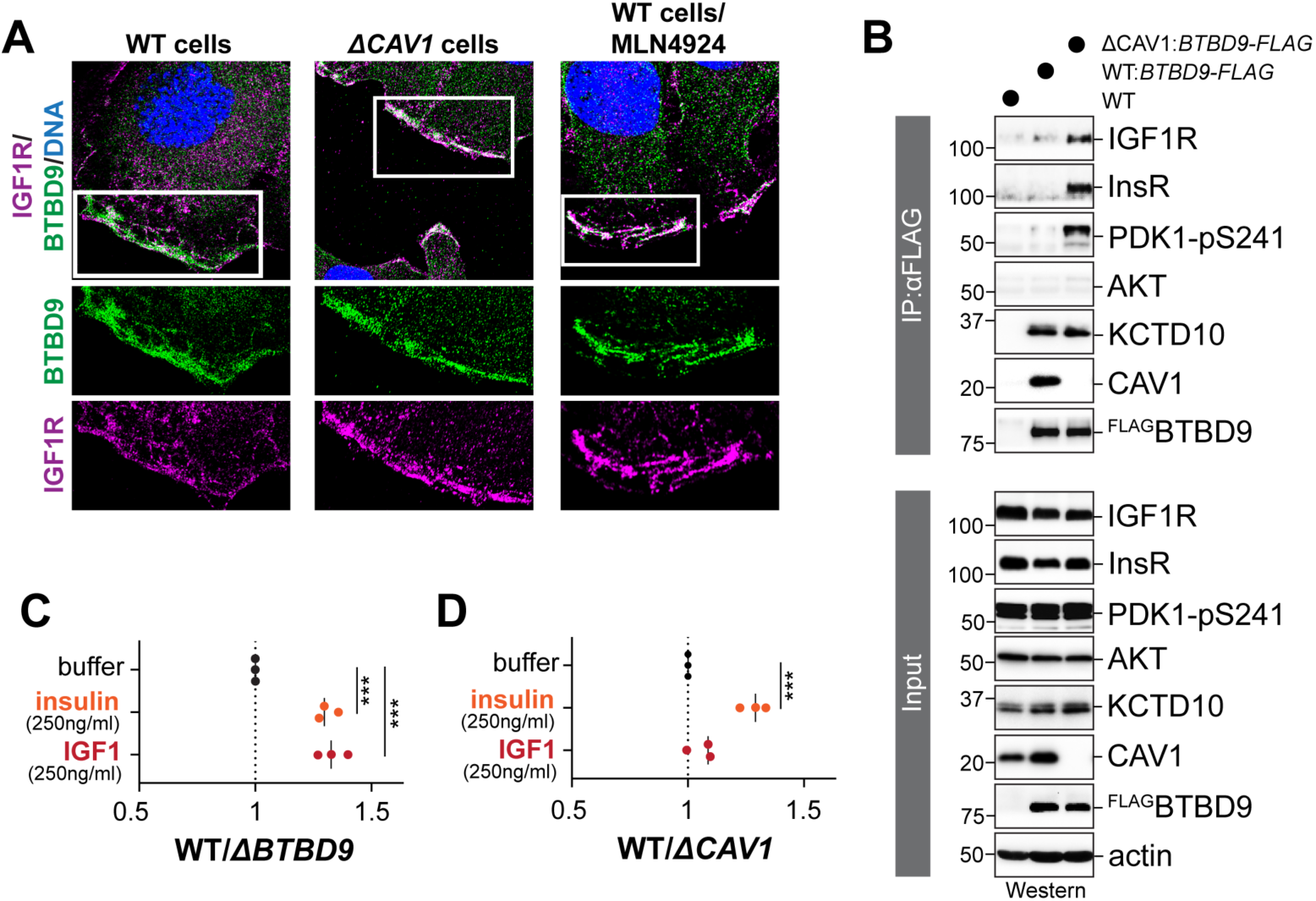
BTBD9 regulates protein dynamics at the plasma membrane. **A.** Localization of endogenous BTBD9 and IGF1-R was analyzed in WT C2C12 myoblasts, *ΔCAV1* myoblasts, or WT myoblasts treated with the CUL3 inhibitor MLN4924. A larger magnification of the insert focusing on plasma membrane-localized proteins is shown below. **B.** Affinity-purification of endogenously FLAG-tagged BTBD9 from either wildtype or *ΔCAV1* myoblasts. Co-precipitating proteins were detected by Western blotting. **C.** Competition assay between wildtype C2C12 myoblasts stably expressing GFP and *ΔBTBD9* myoblasts stably expressing mCherry. Cells were mixed at a 1:1 ratio, briefly serum-starved, and restimulated with either insulin or IGF1. GFP/mCherry-positive cells were detected 24h later by flow cytometry. **D.** Competition assay between wildtype C2C12 myoblasts stably expressing GFP and *ΔCAV1* myoblasts stably expressing mCherry. Cells were mixed at a 1:1 ratio, briefly serum-starved, and restimulated with either insulin or IGF1. GFP/mCherry-positive cells were detected 24h later by flow cytometry.

Consistent with these results, we noted that the association of BTBD9 with insulin or IGF1 receptors was increased in the absence of *CAV1* (**Figure 6B**). BTBD9 bound KCTD10 with similar efficiency under these conditions, suggesting that CAV1 specifically controls interactions of the E3 ligase adaptor with insulin/IGF1 receptors. In addition, *CAV1* deletion increased the presence of active PDK1, the kinase responsible for AKT activation, within BTBD9 complexes (**Figure 6B**). These findings suggested that CAV1 allows CUL3^BTBD9^ to turn over complexes of insulin receptors and effectors at the plasma membrane. We hypothesize that CUL3^BTBD9^-dependent disassembly of receptor complexes at the plasma membrane could allow PDK1 to gain access to AKT, leading to AKT activation and downstream signaling events important for myogenesis.

To provide independent evidence for a role of CUL3^BTBD9^ in insulin- and AKT-signaling, we assessed the response of wildtype or *ΔBTBD9* myoblasts to insulin using a cell competition assay, as reported previously ^40^. We mixed GFP-labeled wildtype and mCherry-labeled *ΔBTBD9* myoblasts and following a brief period of starvation, re-stimulated these cells with either insulin or IGF-1. Following a day of incubation that allows myoblasts to divide once, we measured the relative abundance of each cell type by flow cytometry. Despite the short timeframe of our experiment that allowed for only one round of cell division, wildtype myoblasts outcompeted their *ΔBTBD9* counterparts by almost 50% (**Figure 6C**). We found that *ΔCAV1* myoblasts were similarly impaired in their response to insulin as cells lacking *BTBD9*, but less so towards IGF-1 (**Figure 6D**). Based on these findings, we conclude that CUL3^BTBD9^ and CAV1 collaborate to control insulin signaling in myoblasts as a prerequisite for AKT activation and successful differentiation. Further work is required to delineate the biochemical basis of CUL3^BTBD9^ function in this pathway.

## Discussion

Ubiquitylation has emerged as a central regulator of muscle formation and homeostasis, but E3 ligases that carry out specific modifications in muscle remain to be discovered. Here, we identify CUL3^BTBD9^ as an E3 ligase required for myoblast differentiation. Inactivation of *BTBD9* blocked differentiation prior to induction of myogenin-dependent gene expression. CUL3^BTBD9^ binds and ubiquitylates CAV1, the core component of caveolae that regulate of insulin signaling during myogenesis ^47^, and CAV1 and CUL3^BTBD9^ were both required to ensure insulin-responsiveness and AKT activation in myobasts. Together, these observations suggest that CUL3^BTBD9^ and CAV1 play an important role in regulating insulin signaling to ensure successful myogenesis.

### CUL3^BTBD8^ binds and ubiquitylates CAV1

Shown by the strong effects of its deletion, BTBD9 and by inference the holoenzyme CUL3^BTBD9^ are essential for myoblast differentiation *in vitro*. As most E3 ligases have multiple substrates, the impact of *BTBD9* inactivation is likely due to dysregulation of many targets at the same time. Indeed, different from our study showing that a differentiation block caused by CUL3^KEAP1^ inactivation depletion could be rescued by simultaneous deletion of its degradation target NRF2 ^19^, we were unable to identify a single substrate that was responsible for the effects of BTBD9 loss on myogenesis.

Mass spectrometry analyses showed that BTBD9 abundantly binds CAV1, the major component of membrane domains referred to as caveolae. Caveolae are important for myogenesis and insulin signaling ^48–50^. CUL3^BTBD9^ decorates CAV1 with ubiquitin oligomers that range in size from one to three subunits, and as expected, such conjugates did not drive CAV1 degradation. Instead, the similar effects of *BTBD9* and *CAV1* deletion on AKT function suggested that ubiquitylation activates a CAV1 function required for insulin-dependent AKT stimulation. Our findings are in line with other CUL3-family E3 ligases predominantly catalyzing monoubiquitylation to regulate protein interactions and function ^51–55^. BTBD9 recognizes the CAV1 scaffolding domain that recruits multiple signaling proteins ^28^, and CUL3^BTBD9^ most abundantly modifies Lys residues in the neighboring CAV1 amino-terminus that is also subject to regulatory phosphorylation ^56^. Mass spectrometry analyses revealed multiple CAV1 interactions that are lost in the absence of *BTBD9*, and some of these binders, such as GNAI3, LAT1, or JPH1, are required for myogenesis. It is possible that ubiquitylation through CUL3^BTBD9^ increases the affinity of CAV1 to such partners. Addressing this question requires reconstitution of CAV1 modification *in vitro*, which we have not yet accomplished potentially due to a potential requirement for posttranslational modifications or additional ubiquitylation factors that remain to be discovered.

Prior to our work, CAV1 had been found to be the target of other E3 ligases. ZNRF1 modifies CAV1 with ubiquitin chains that mediate proteasomal degradation and thereby regulate Toll-like receptor dependent immune-signaling ^57^. ZNRF1 decorates the same K39 residue in CAV1 as CUL3^BTBD9^, and it restricts activation of the same kinase, AKT, by limiting CAV1 accumulation. These observations suggest that AKT is a central effector of CUL3^BTBD9^-dependent ubiquitylation. The ER-associated HRD1 can ubiquitylate CAV1 during pathological conditions, such as ischemic stroke ^58^. Modification of CAV1 by an unknown E3 ligase also triggers its sorting to lysosomes for degradation independently of proteasomes ^59^. Interestingly, this ubiquitylation also occurs at amino-terminal residues of CAV1, and it recruits the p97/VCP segregase that is known to remodel protein complexes. Thus, the amino-terminus of CAV1 is a hotspot for ubiquitin-dependent regulation, the consequences of which depend on the responsible E3 ligase and its inherent ubiquitin chain topology.

### CUL3^BTBD9^ regulates insulin signaling

While studying CAV1 interactions in myoblasts, we noted that *BTBD9* deletion caused a strong increase in the expression of the *CAV1* gene. CAV1 expression is regulated by FOXO transcription factors that are inhibited by AKT ^44^, and multiple other FOXO target genes were also upregulated by the absence of *BTBD9*. The effects of *BTBD9* deletion on gene expression were most prominent after myoblasts had been transferred to differentiation medium that contains low concentrations of growth factors required to sustain AKT activation. Together, these findings suggested that CUL3^BTBD9^ plays a role in growth factor-dependent activation of AKT.

Consistent with this hypothesis, we found that BTBD9 localizes to plasma membranes. The inactivation of CAV1 resulted in a stronger co-localization of BTBD9 with IGF1-receptors that translate IGF1 recognition into AKT activation. We could not detect insulin receptor in myoblasts by microscopy, but assume that BTBD9 also co-localizes with IR more efficiently if CAV1 has been deleted. Mirroring the changes in localization, deletion of *CAV1* increased the association of BTBD9 with insulin and IGF1 receptors, and these complexes also contained active PDK1 kinase that usually would stimulate downstream AKT. We therefore propose that CUL3^BTBD9^-dependent ubiquitylation of CAV1 remodels insulin and IGF1 receptor complexes at the plasma membrane to release PDK1 as a prerequisite for subsequent AKT activation.

How CUL3^BTBD9^ accomplishes such regulation is still unclear. In one model, CUL3^BTBD9^ catalyzes CAV1 ubiquitylation to dissociate it from receptors and thereby release proteins of the same complexes, such as PDK1. Consistent with this notion, ubiquitylation on K39 of CAV1 has previously been reported to recruit the p97/VCP disaggregase that dismantles protein complexes in a ubiquitin-dependent manner ^59^. Alternatively, CAV1 ubiquitylation might increase its affinity to a receptor complex component and competitively release proteins, such as PDK1. This possibility would be consistent with a compensatory transcriptional upregulation of CAV1, as well as PI3K, in cells lacking *BTBD9*. While more work is needed to dissect the biochemical basis for CUL3^BTBD9^ and CAV1 activity, our work suggests that a ubiquitin-dependent reaction occurring the plasma membrane plays important roles in myogenic signaling. This illustrates how ubiquitylation controls membrane-associated signaling during differentiation, opening the door to mechanistic studies that could reveal fine-tuning of insulin signaling in development and disease.

## Supplemental Figures and Legends

**Figure S1:**
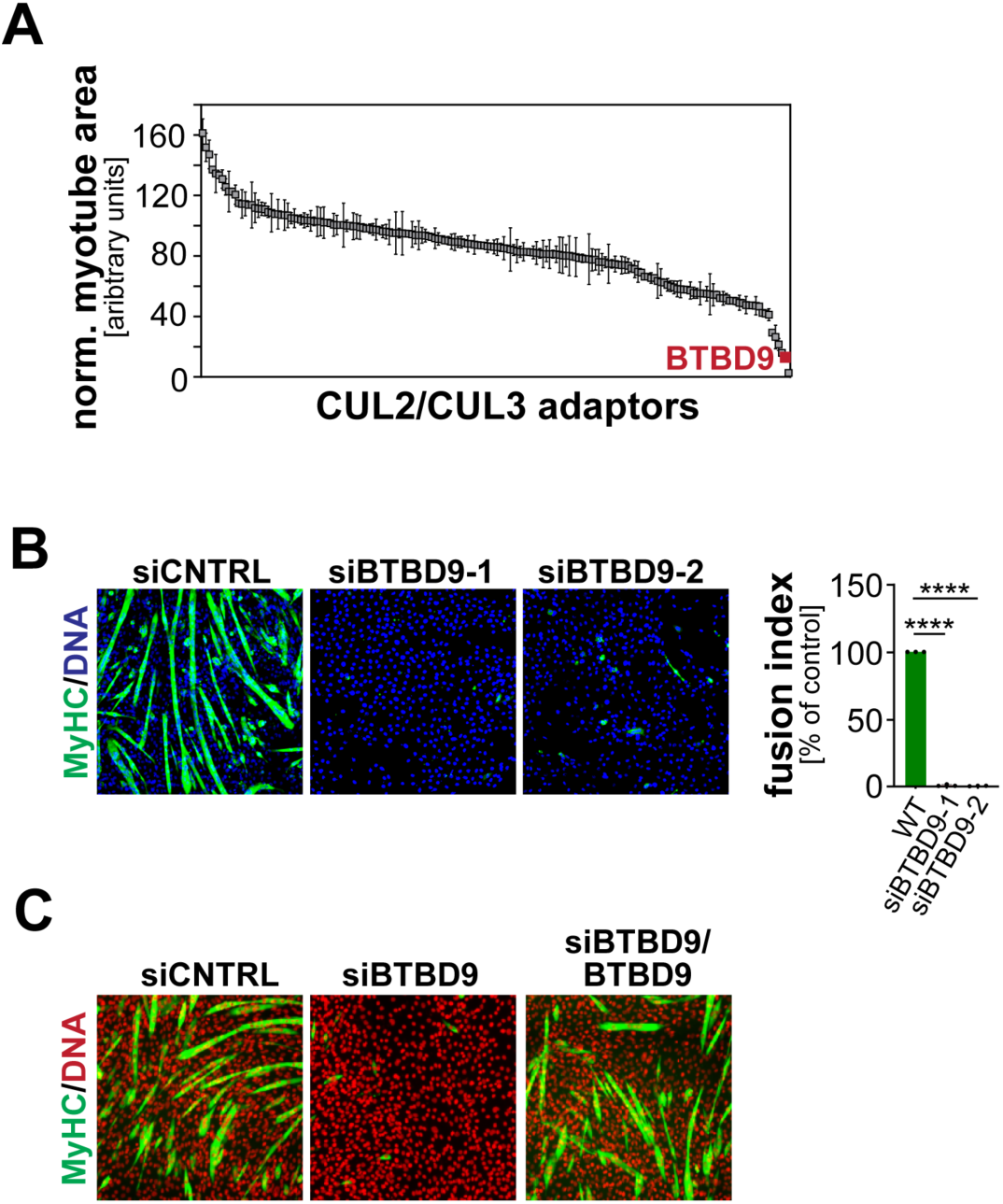
BTBD9 is required for C2C12 differentiation. **A.** BTBD9 depletion inhibits C2C12 myogenesis, as found in a previously reported genetic screen. The screen data was derived from ref. ^19^. **B.** Depletion of BTBD9 with two independent siRNAs blocks C2C12 differentiation, as detected by the lack of MyHC expression. Quantification of three independent experiments is shown on the right. **C.** Differentiation of siBTBD9-treated myoblasts could be restored by expression of an siRNA-resistant version of BTBD9. Differentiation was followed by monitoring expression of MyHC by microscopy.

**Figure S2:**
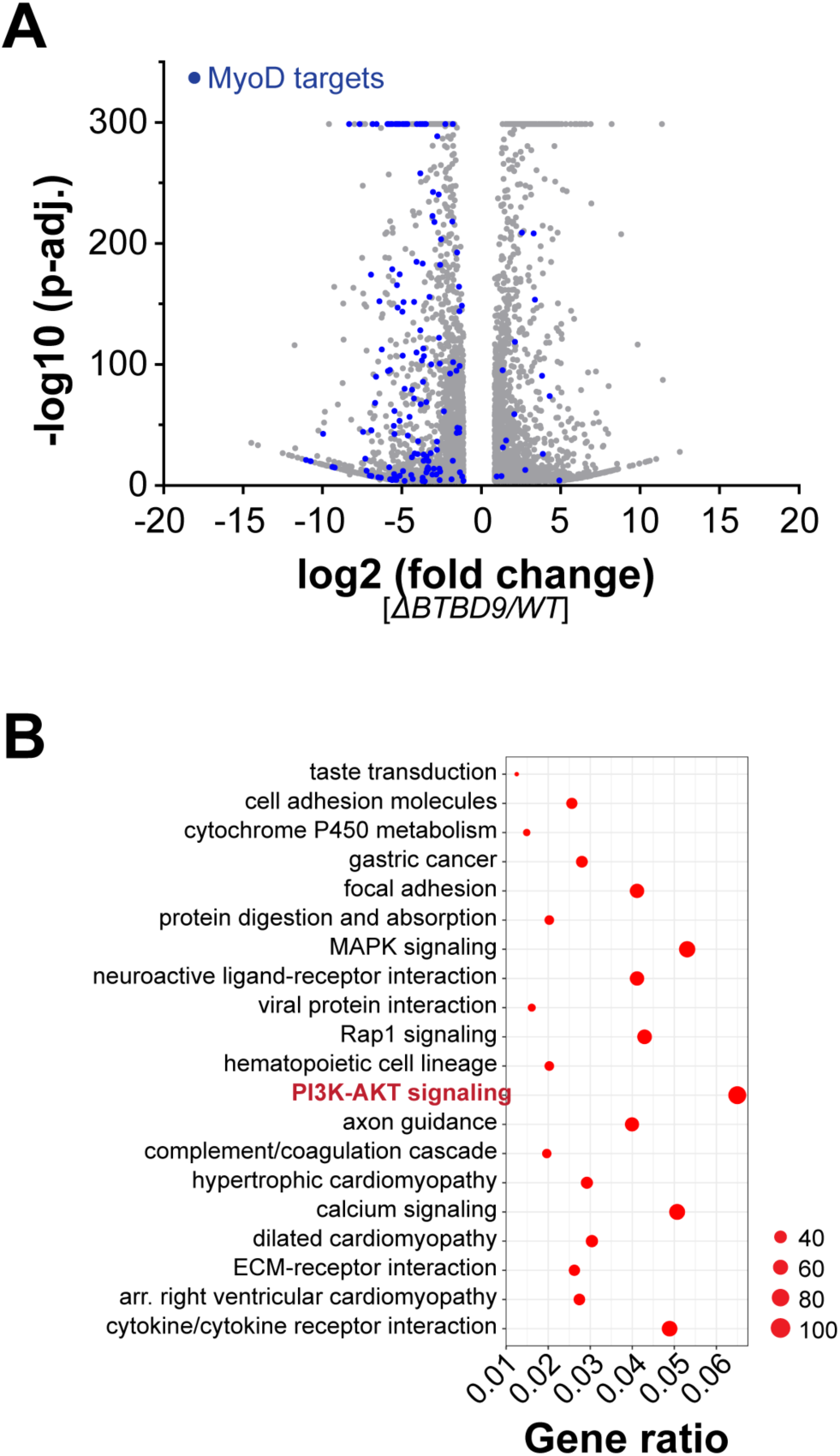
Gene expression changes caused by *BTBD9* deletion. **A.** Wildtype or *ΔBTBD9* C2C12 myoblasts were transferred to differentiation medium. 2d later, gene expression was analyzed by RNA-sequencing (*blue:* targets of the muscle transcription factor MyoD). **B.** Go term analysis of pathways dysregulated by *BTBD9* deletion in C2C12 myoblasts subjected to 2d differentiation.

**Figure S3:**
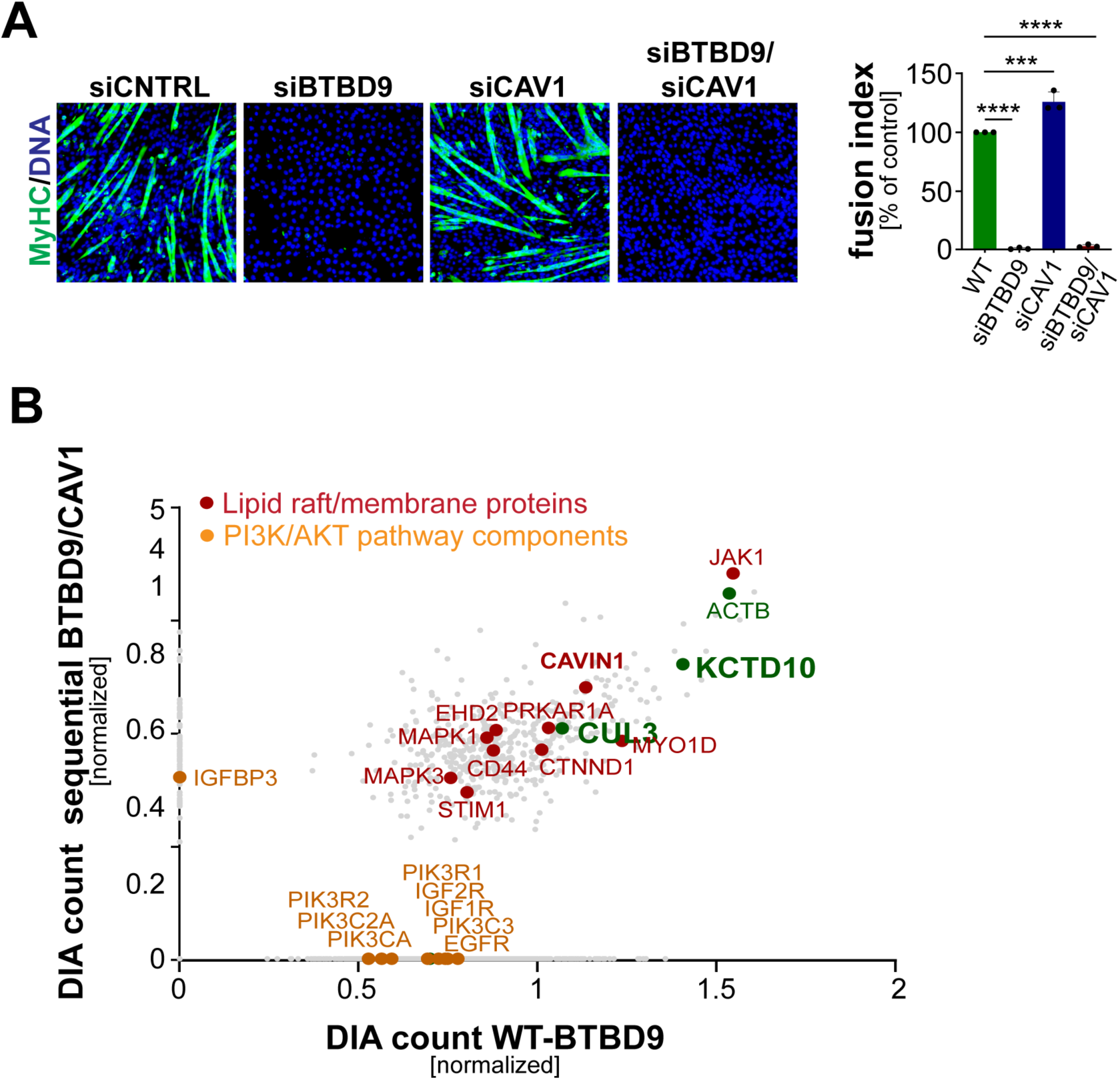
A. Characterization of CAV1 function and interactions. **A.** C2C12 myoblasts were treated with control siRNAs or siRNAs targeting BTBD9, CAV1, or both. Cells were transferred to differentiation medium, and myotube formation was followed by immunofluorescence microscopy against MyHC. Quantification of three independent experiments is shown on the right. **B.** Sequential affinity-purification of BTBD9 and CAV1 compared to single BTBD9 immunoprecipitation, as analyzed by mass spectrometry. KCTD10, Cavin1, and CUL3 are found in both purifications, while many components of the PI3K pathways are only found in BTBD9-purifications.

**Figure S4:**
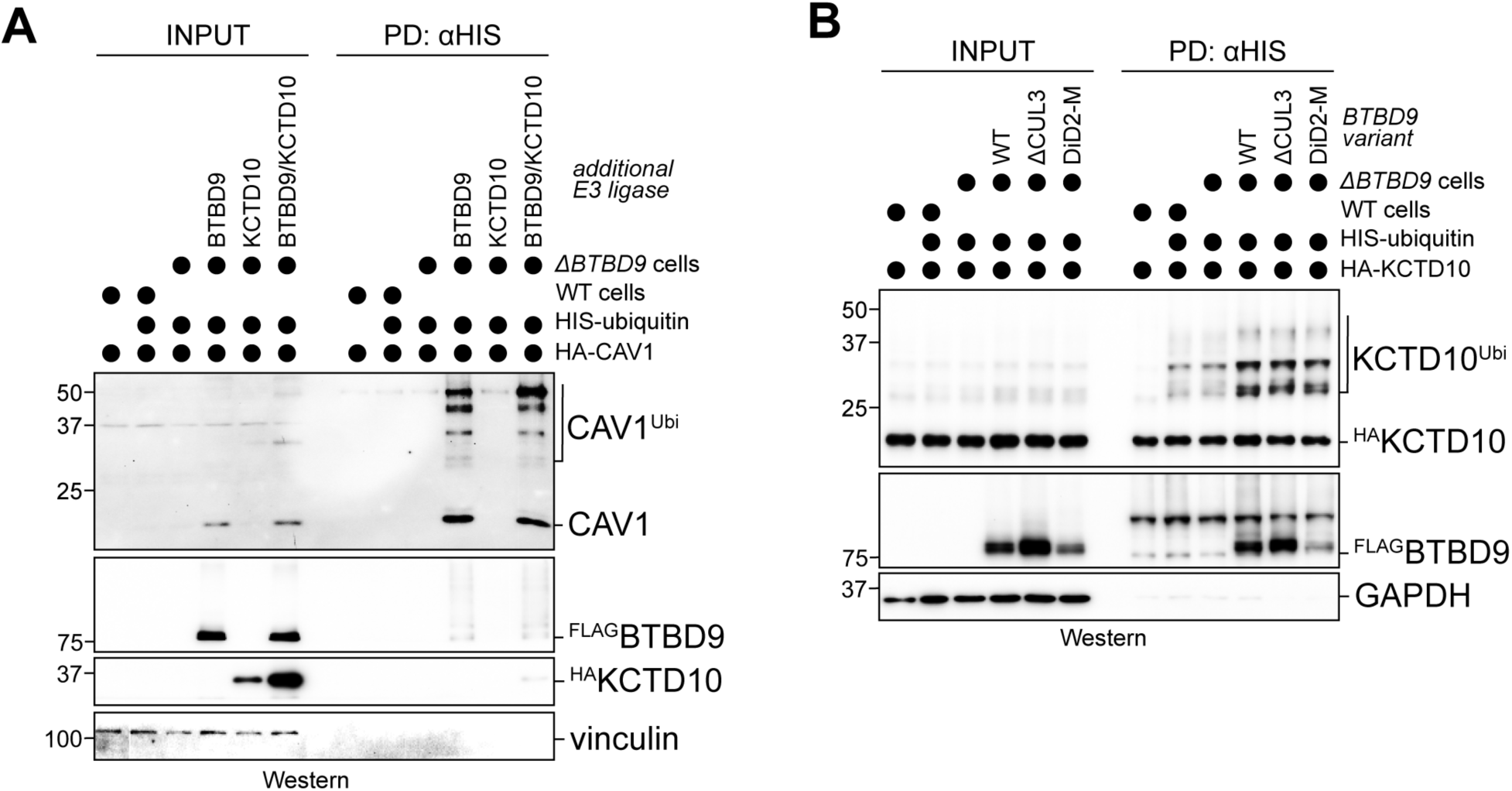
Analysis of cellular ubiquitylation of CAV1 and KCTD10 upon BTBD9 over-expression. **A.** Denaturing NiNTA-purification of His-ubiquitin conjugates from either WT or *ΔBTBD9* myoblasts shows that overexpression of BTBD9 resulted in modification of CAV1 with up to three ubiquitin molecules, as detected by Western blotting. As indicated, KCTD10 was expressed either by itself or in combination with BTBD9. **B.** Denaturing NiNTA-purification of His-ubiquitin conjugates from either WT or *ΔBTBD9* myoblasts showing that ubiquitylation of KCTD10 occurs independently of whether BTBD9 can bind CUL3 or substrate, as detected by Western blotting.

## Material and Methods

### Mammalian cell culture

HEK293T and HeLa cell lines were purchased from the UC Berkeley Cell Culture Facility, where cells are authenticated by short tandem repeat analysis, and C2C12 myoblasts were purchased from ATCC (CRL-1772). All cell lines were maintained in Dulbecco’s modified Eagle’s medium (DMEM) + GlutaMAX (GIBCO, 51985091) supplemented with 10% fetal bovine serum (FBS) (VWR, 97068-107). All cells were regularly tested and found negative for mycoplasma.

To differentiate C2C12 myoblasts, the growth medium was changed from DMEM + 10% FBS to DMEM + 2% horse serum. Cells were allowed to differentiate for 3-4 days, and media was changed daily.

Lipofectamine RNAiMAXX (Thermo Fisher Scientific, 13778030) was used for all siRNA transfections according to manufacturer’s instructions. Final concentration of siRNA ranged from 30 to 60nM, depending on the experiment. For siRNA-depletions in myoblast differentiation experiments, C2C12s were seeded at 40,000 cells per well in 12-well plates and transfected with siRNAs two days prior to switching to low serum media.

Plasmid transfection of HEK293Ts were performed using polyethylenimine (PEI; Polysciences, 23966-1) at a ratio of 1:6 µg DNA to µg/mL PEI or Lipofectamine 3000 Transfection Reagent (Thermo Fisher Scientific, L3000015) according to the manufacturer’s protocol.

### Plasmids

All plasmids used in this study are listed in Supplementary Table 1. Site-directed mutagenesis was performed using the quick change method. All other cloning was completed by Gibson assembly using HIFI DNA Assembly master mix (NEB, E2621L). PrimeSTAR GXL DNA Polymerase (Takara, R050A) was used for all PCR reactions. Genes of interest were amplified from HEK293T or C2C12 cDNA produced using Protoscript II First Strand cDNA Synthesis Kit (NEB, E6560S).

### Virus production and infection

Lentivirus was produced by co-transfection of HEK293T cells with pLVX or pINDUCER20 plasmids and lentiviral packaging plasmids using PEI. Media from transfected cells was collected after 48 and 72h, pooled, and spun down. Lenti-X Concentrator (Takara Bio, 631262) was used to concentrate viral particles according to the manufacturer’s instructions. Once concentrated, virus-containing pellets were resuspended in 2mL DMEM + 10% FBS, aliquoted, and stored at - 80°C.

For infections, 100,000 HeLa cells or 50,000 C2C12s were seeded in 12-well plates with 8ug/mL polybrene (Sigma-Aldrich, TR-1003) and 80-125uL concentrated virus and centrifuged at 1,000 x g for 45min at 30°C. After 24h, cells were transferred to 10-cm plates and drug-selected with 1mg/mL geneticin (GIBCO, 10131-035), 0.5µg/mL puromycin (Sigma-Aldrich, P8833) or 10µg/mL blasticidin (Thermo Fisher Scientific, A1113903).

### Generation of CRISPR-cas9 genome edited cell lines

BTBD9 and CAV1 deletions in HEK293T, HeLas, and C2C12s were generated using predesigned Gene Knockout Kits purchased from EditCo Bio, Inc. Cells were nucleofected with RNP complexes using Cell Line 4-D Nucleofector Kits purchased from Lonza according to manufacturer’s instructions. After bulk editing was confirmed by PCR, cells were seeded in 96-well plates at ∼1 cell per well for single-cell selection. Individual clones were checked by PCR for editing and then validated by sequencing, western blotting or IP-mass spectrometry.

### Western blotting from whole cell lysates

Cells for western blotting were trypsin digested, quenched with DMEM + 10% FBS, spun down, washed with 1X DPBS, snap-frozen, and stored at -80°C. Samples were resuspended in swelling buffer (20mM HEPES pH 7.5, 5mM KCl, 1.5mM MgCl2, 150mM NaCl, 0.1% NP-40, Benzonase Nuclease (EMD Millipore, 70746-4), Roche complete EDTA-free protease inhibitor cocktail (Sigma-Aldrich, 11873580001) and PhosSTOP phosphatase inhibitor cocktail (Roche, 4906837001)), incubated on ice for 20 min and then combined with 2X urea sample buffer. Samples were then normalized for protein concentration using Pierce 660 Protein Reagent (Thermo Scientific, 22660) plus IDCR reagent (Thermo Scientific, 22663). Samples were separated by SDS-PAGE, transferred to nitrocellulose membranes and blocked in 1X PBS + 0.1% Tween-20 (Sigma, P7949) for 30min at room temperature. Primary antibodies were added to membranes for overnight incubation at 4°C. Membranes were incubated with HRP-conjugated secondary antibodies at 1:5000 dilution for 1h at room temperature, treated with Immobilon Western Chemiluminescent Substrate (Millipore, WBKLS05000) according to manufacturer’s instructions, and imaged with a ProteinSimple FluorChem M device. Membranes were washed three times with 1X PBST between and after each antibody incubation.

For serum-starvation plus insulin stimulation experiments, C2C12s were seeded in 6-well plates at 100,000 cells per well. After 48h, cells were switched to serum-free DMEM for 2.5h, treated with 200ng/mL (∼34nM) recombinant human insulin (Selleckchem, S6955) for 20min, harvested, and stored at -80°C until western blotting.

### Antibodies

All antibodies and their dilutions used in this study are listed in Supplementary Table 2.

### siRNAs

The following siRNAs were used in this study::

ON-TARGETplus mouse Btbd9 siRNA #1 (GAACGAAAGUGUCCUGCAA)

ON-TARGETplus mouse Btbd9 siRNA #2 (GCGCUGAAGGAAUCUACAA)

ON-TARGETplus mouse Cav1 siRNA #4 (GUCCAUACCUUCUGCGAUC)

### Small-scale immunoprecipitation

HEK293T cells were seeded at 2E6 cells per 10-cm plate and transiently transfected with plasmid DNA the following day using PEI. After 48hrs, cells were harvested, washed with 1X PBS, pelleted and snap-frozen. Pellets were resuspended in lysis buffer (30mM TrisHCl pH 7.6, 150mM NaCl, 0.5% NP-40, with Roche complete EDTA-free protease inhibitor cocktail ((Sigma-Aldrich, 11873580001) and PhosSTOP phosphatase inhibitor cocktail (Roche, 4906837001), incubated for 30min at 4°C, and centrifuged at 15,000 rpm for 10min at 4°C. Supernatants were collected and normalized to volume and protein concentration using Pierce 660 Protein Reagent (Thermo Scientific, 22660) plus IDCR reagent (Thermo Scientific, 22663). Samples were then incubated with anti-FLAG M2 affinity resin (Sigma, A2220) for 3h at 4°C. Beads were washed 4 times with wash buffer (30mM TrisHCl pH 7.6, 150mM NaCl, 0.5% NP-40). After the final wash, supernatants were completely removed using gel loading tips, and 2X urea sample buffer was added. Samples were then separated by SDS-PAGE and transferred to nitrocellulose membranes for western blotting.

For endogenous BTBD9-3xFLAG coIPs, four 15-cm plates per condition were grown to ∼90% confluency, scraped into 10mL 1X DPBS and pooled, spun down, washed with 1X DPBS and snap frozen. Immunoprecipitation was completed as described above.

### Immunoprecipitation and mass spectrometry

^FLAG^BTBD9, ^FLAG^BTBD9^ΔCUL3^ and ^FLAG^BTBD9^NEV/AKD^ mass spectrometry experiments were performed from 15 15-cm plates of lentiviral stable HeLa cell lines with dox-inducible expression. Cells were seeded at 4e6 per 15-cm plate, induced with 1ug/mL doxycycline for 48h, and harvested as described above. Immunoprecipitation was performed following the same protocol as small-scale immunoprecipitation, but with the following change. After the final wash, the beads were further washed with 1X PBS to remove NP-40 and snap-frozen in a small volume of 1X PBS.

Further sample processing was performed at UC San Diego Proteomics Facility as described previously ^40^.

For the CAV1^HA^ mass spectrometry experiment, 15 15-cm plates of wild-type and ΔBTBD9 HeLa cells stably expressing CAV1^HA^ were seeded, doxycycline-induced, and immunoprecipitated as described for the ^FLAG^BTBD9 mass spectrometry experiments. Three technical replicates were used to generate volcano plots.

### Immunofluorescence microscopy

C2C12 cells were seeded at 5,000 to 10,000 cells/mL on cover slips in 12-well plates. After 48h, cells were treated with DMSO or 1µM MLN-4924 (Cayman Chemical, 15217) for 18h. Cells were then fixed in 1X PBS + 4% paraformaldehyde for 15 min, permeabilized in 1X PBS + 0.1% Triton X-100 for 15 min, and blocked in 1X PBS + 10% FBS for 30min. For antibody staining, cells were incubated for 3h in 1X PBS + 10% FBS containing anti-IGF1R (Abcam, AB182408) and anti-FLAG (Sigma, F1804) at 1:250 dilutions. Cells were washed with 1X PBS + 10% FBS and then incubated with goat anti-mouse AlexaFlour 488 (1:500; Thermo Fisher, A32723), goat anti-rabbit AlexaFluor 647 (1:500; Thermo Fisher, A32733), and Hoechst 33342 (1:3000; AnaSpec, 83218) for 1h. Finally, cells were washed with 1X PBS and mounted on glass slides with ProLong Gold antifade reagent (Thermo Fisher, P36930). Imaging was performed on a Zeiss LSM900 Airyscan 2 using Airyscan settings at 63x.

### Myogenesis functional assays

C2C12 cells were seeded at 40,000 cell per well in 12-well plates, grown to ∼90% confluency and differentiated. Fixation, permeabilization, and blocking was performed as described in the immunofluorescence microscopy section. Depending on the experiment, cells were stained with the primary antibody anti-myosin heavy chain (DHSB; MF20) at 1:50, anti-myogenin (DHSB; F5G) at 1:50, or anti-CAV1 (Cell Signaling, 3267S) at 1:200 for 3h. After washing with 1X PBS, cells were incubated with secondary antibodies goat anti-mouse AlexaFluor 488 (Thermo Fisher, A32723) or goat anti-rabbit AlexaFluor 488 (Thermo Fisher, A32731) at 1:500 and Hoechst 33342 (AnaSpec, 83218) at 1:3000 for 1h at room temperature. Samples were imaged using a Perkin Elmer Opera Phenix, and data was analyzed using Harmony image analysis software.

### Fusion index analysis

To quantify myogenesis phenotypes in myogenesis functional experiments, the fusion index parameter was calculated. It is defined as the number of nuclei within MyHC+ cells (containing ≥ 3 nuclei) divided by the total number of nuclei. Data are presented as mean ± standard deviation.

### His-Ub pulldowns

293T parental or 293T *ΔBTBD9* cells were seeded at 2e6 cells per 10-cm plate. The following day, cells were transiently transfected with pCS2 plasmids of CAV1-HA, 6xHis-ubiquitin, and 3xFLAG-BTBD9 using PEI at ratio of 6:1 (ug PEI to ug plasmid DNA) for 48h. Cells were then harvested, washed with 1X PBS and snap-frozen.

Cell pellets were resuspended in urea lysis buffer (8M urea, 300mM NaCl, 0.5% NP-40, 50mM Tris-HCl pH 7.6) made from fresh 10M urea solution and incubated at room temperature for 20min. Lysates were sonicated with microtip attached at 20 Amp for a total of 10 sec (1 sec on, 1 sec off) and then centrifuged at 15,000 rpm for 15min. Supernatants were transferred to fresh eppendorf tubes and normalized for protein concentration and volume. An equal amount of equilibrated Ni-NTA agarose (QIAGEN, 30210) was added to each sample, followed by a 1h incubation at room temperature on a rotator. Ni-NTA samples were then spun down and washed three times with urea lysis buffer and twice with wash buffer (8M urea, 300mM NaCl, 50mM Tris-HCl pH 7.6). Finally, supernatants were removed, and 50uL elution buffer (2X urea sample buffer + 200mM imidazole) was added. Samples were either immediately analyzed by western blotting or stored at -20°C for later analysis.

### Membrane fractionation

C2C12 cells were seeded in three 15-cm plates per condition and grown to ∼90% confluency. Cells were then serum-starved for 2h, stimulated with 200ng/mL insulin (Selleckchem, S6955) for 20min, and harvested as described previously. Pellets were resuspended in lysis buffer (20mM HEPES, 7.4, 250mM sorbitol, 1mM EDTA with protease and phosphatase inhibitor cocktails), passed 12 times through a 22 gauge syringe for lysis, and centrifuged at 1,000 x g for 10min at 4°C. Supernatants were collected, and the membrane were pelleted by ultracentrifugation at 100,000 x g for 30min at 4°C. The supernatants were removed for the cytosolic fractions, and the pelleted membrane fractions were then resuspended in lysis buffer. SDS-PAGE samples were taken from the 1,000 x g centrifugation supernatants, the 100,000 x g centrifugation supernatants, and the resuspended membrane pellets for inputs, cytosolic fractions, and membrane fractions, respectively.

### Growth competition assays

C2C12 parental cells were infected with pLVX-EF1ɑ-mCherry-Blast, and C2C12 *ΔBTBD9* or *ΔCAV1* cells were infected with pLVX-EF1ɑ-GFP-Blast. mCherry- and GFP-expressing cells were then mixed 1:1 and seeded in 12-well plates at 25,000 cells per well. After 48hr, cells were switched to serum-free DMEM and 250ng/mL recombinant human insulin (Selleckchem, S6955) or 250ng/mL IGF1 (Millipore Sigma, I3769) was added. After 24h, the cells were harvested and analyzed by flow cytometry using a BD LSR Fortessa flow cytometer. Cell ratios were calculated using Flowjo software.

### Reporter stability assays

WT and *ΔBTBD9* HEK293T cells were seeded in 6-well plates at 3e5 cells per well. The following day, cells were transiently transfected with pCS2-CAV1^FL^-mEmerald-IRES-mCherry or pCS2-CAV1^1-102^-mEmerald-IRES-mCherry using Lipofectamine 3000 according to the manufacturer’s instructions. After 24h, mEmerald and mCherry signals were measured using a BD LSR Fortessa flow cytometer. Using Flowjo software, the mEmerald to mCherry ratio was calculated as a readout for reporter stability.

### RNA-Seq

Wild-type and *ΔBTBD9* C2C12 cells were seeded in 6-well plates and harvest at day 0 and day 2 of differentiation with three biological replicates per condition. RNA was extracted using the NucleoSpin RNA kit (Macherey-Nagel, 740955). RNA samples were sent to Novogene, where library prep, sequencing, quality control and analysis were performed. Transcription factor enrichment analysis was done using the Enrichr online tool ^60–62^.

### RT-qPCR

RNA extraction was performed as described above. To generate cDNAs, 1ug of RNA was reverse transcribed using the Protoscript II First Strand cDNA Synthesis kit (NEB, E6560S). RT-qPCR was performed in 384-well plates using the KAPA SYBR Fast Universal qPCR kit (Roche, KK4602) and on a LightCycler 480 II instrument (Roche). Finally, the ΔΔCt method was used to calculate relative expression levels. The qPCR primers used in this study are listed in Supplementary Table 3.

## QUANTIFICATION AND STATISTICAL ANALYSIS

The quantifications in this study are presented as the mean ± standard deviation (SD). All myogenesis experiments, RT-qPCR, RNA-seq, and FACS-based growth competitions were performed using three biological replicates. GraphPad Prism was used to analyze for statistical significance using either the one sample t-test or one-way ANOVA to compare the desired conditions (* p ≤ 0.05, ** p ≤ 0.01, *** p ≤ 0.001, **** p ≤ 0.0001).

## Acknowledgements

We thank all members of our laboratory for enthusiastic support and many suggestions for this work. We thank Dr. Majid Ghassemian and the UCSD Biomolecular and proteomics mass spectrometry facility for assistance in running the mass spec samples. MR is an Investigator of the Howard Hughes Medical Institute.

## Conflict of interest statement

MR is co-founder and SAB member of Nurix Therapeutics; co-founder, consultant and SAB member of Lyterian Therapeutics; co-founder and consultant of Zenith Therapeutics; co-founder of Reina Therapeutics; and iPartner at The Column Group. All other authors declare no conflicts of interest.

